# AB668, a novel highly selective protein kinase CK2 inhibitor with a distinct anti-tumor mechanism as compared to CX-4945 and SGC-CK2-1

**DOI:** 10.1101/2022.12.16.520736

**Authors:** Alexandre Bancet, Rita Frem, Florian Jeanneret, Angélique Mularoni, Pauline Bazelle, Caroline Roelants, Jean-Guy Delcros, Jean-François Guichou, Catherine Pillet, Isabelle Coste, Toufic Renno, Christophe Battail, Claude Cochet, Thierry Lomberget, Odile Filhol, Isabelle Krimm

**Affiliations:** Univ Lyon, Université Claude Bernard Lyon 1, INSERM 1052, CNRS 5286, Centre Léon Bérard, Centre de recherche en cancérologie de Lyon, Institut Convergence Plascan, Team « Small Molecules for Biological Targets », Lyon, 69373, France; Univ Lyon, Université Claude Bernard Lyon 1, INSERM 1052, CNRS 5286, Centre Léon Bérard, Centre de recherche en cancérologie de Lyon, Institut Convergence Plascan, Team Targeting non-canonical protein functions in cancer, Lyon, 69373, France; Univ. Grenoble Alpes, INSERM 1292, CEA, UMR Biosanté, Grenoble, 38000, France; Centre de Biologie Structurale, CNRS, INSERM, Univ. Montpellier, Montpellier 34090, France; Univ Lyon, Université Claude Bernard Lyon 1, CNRS UMR 5246, Institut de Chimie et Biochimie Moléculaires et Supramoléculaires (ICBMS), COSSBA Team, Faculté de Pharmacie-ISPB, 8 Avenue Rockefeller, FR-69373 Lyon Cedex 08, France

## Abstract

Although the involvement of protein kinase CK2 in cancer is well-documented, there is a need for selective CK2 inhibitors suitable for investigating CK2 specific roles in cancer-related biological pathways and further explore its therapeutic potential. Here we have discovered AB668, a new bivalent inhibitor that binds both at the ATP site and an allosteric αD pocket unique to CK2. The molecule inhibits CK2 activity with an outstanding selectivity over other kinases. Using caspase activation assay, live-cell imaging and transcriptomic analysis, we have compared the effects of this bivalent inhibitor to the non-selective ATP-competitive inhibitor CX-4945 that reached clinic and to the selective ATP-competitive SGC-CK2-1 molecule. Our results show that in contrast to CX-4945 or SGC-CK2-1, AB668 has a distinct mechanism of action regarding its anti-cancer activity, inducing apoptotic cell death and stimulating distinct biological pathways in several cancer cell lines while sparing healthy cells. Our data suggest that targeting a cryptic CK2 αD pocket validates an allosteric approach to targeting CK2 and provides a starting point for creating drug-like CK2 inhibitors for aggressive cancers.

## Introduction

CK2 is a constitutively active protein kinase ubiquitously expressed in eukaryotes and particularly well conserved among species. CK2 phosphorylates serine or threonine residues within an acidic context (S/TXXD/E/pS/pT/pY) and is responsible for the generation of a large proportion of the human phosphoproteome (Franchin et al., 2015). One of the distinctive features of CK2 is to exist in cells as a catalytic subunit CK2α and as a holoenzyme CK2α_2_β_2_, which consists of two catalytic subunits CK2α that interact with a dimer of two regulatory subunits CK2β (Niefind 2001). Although the so-called regulatory CK2β subunits are not essential per se for CK2α kinase activity as the catalytic subunit is constitutively active (Niefind et al., 2009), several reports showed that CK2β influences significantly CK2 substrate preference as well as CK2α localization (Núñez de Villavicencio-Díaz et al., 2015; Bibby et al., 2005). Up to now, several hundreds of proteins have been shown to be CK2 substrates in cells (Meggio and Pinna et al., 2003; Bian 2016; Borgo et al., 2017). CK2 is involved in important physiological functions, such as embryonic development, differentiation, immunity, cell survival, epithelial homeostasis and circadian rhythms. CK2 is also implicated in numerous human diseases such as cancer, neurodegenerative diseases, viral and parasite infections, cystic fibrosis, psychiatric disorders, diabetes, inflammatory and cardiovascular diseases (Borgo et al., 2021; Dominguez et al., 2009). In cancer, CK2 promotes cell proliferation and survival (Chua et al., 2017a; Trembley et al., 2009). CK2 might also be involved in immune cell development and function in cancer (Husain et al., 2021), in cancer metabolism (Silva-Pavez et al., 2020) as well as in antitumor drug resistance (Borgo et al., 2019). For example, proliferation, migration, invasion, and survival of cholangiocarcinoma cells exposed to cytostatic drugs are markedly reduced when cells are depleted in CK2α subunit (Di Maira et al., 2019). Globally, CK2 is overexpressed in a very large number of human tumors (e.g., breast, ovarian, prostate, lung, colon, kidney, skin and pancreatic cancers) and its over-expression correlates with poor prognosis (Laramas et al., 2007; Chua et al., 2017b; Roelants et al., 2015). The higher sensitivity of cancer cells to CK2 inhibition, as compared to their healthy counterparts, led to the hypothesis of a “non-oncogenic” CK2 addiction of cancer cells (Luo et al., 2009; Ruzzene et al., 2010). Compiling evidence from the literature suggests that CK2 modulates all hallmarks of cancer (Firnau et al., 2022). Consequently, CK2 is considered as a “master regulator” and a promising therapeutic target to treat different human tumors (Roffey et al., 2021).

Many CK2 inhibitors that target the CK2 catalytic site have been proposed (Atkinson et al., 2021). The most advanced molecule silmitasertib (CX-4945) inhibits CK2α catalytic activity with a K_i_ of 0.38 nM (Pierre et al., 2011) and the compound has entered several clinical trials. Notably, silmitasertib received an orphan drug designation for the treatment of cholangiocarcinoma in 2016, medulloblastoma in 2020, recurrent Sonic Hedgehog-driven medulloblastoma in 2021 and biliary tract cancer in 2022 (Lee et al., 2022). Although described as highly selective, CX-4945 inhibits several kinases with nanomolar IC50 values (CLK1, CLK2, CLK3, DYRK1A, DYRK1B, DAPK3, HIPK3…) (Pierre et al., 2011). For example, CX-4945 was reported to regulate splicing in mammalian cells in a CK2-independent manner through the inhibition of Clk1, Clk2 and Clk3 (Kim et al., 2016). Therefore, despite its therapeutic efficacy, CX-4945 cannot be used to probe the cellular functions of CK2. Recently, a newly developed ATP-competitive CK2 inhibitor, SGC-CK2-1, was reported (Wells et al., 2021). SGC-CK2-1 (IC_50_ 2.3 nM on CK2α) is much more selective than CX-4945 and was consequently described as a genuine chemical probe to assess the consequences of the pharmacological inhibition of CK2 kinase activity (Wells et al., 2021). When tested on a panel of 140 cancer cell lines, SGC-CK2-1 reduced cell growth in blood, head/neck, brain, breast, skin, stomach and duodenum cell lines with a micromolar range efficacy. This poor antiproliferative effect led the authors to question the inhibition of CK2 as a strategy for cancer therapy (Wells et al., 2021; Licciardello et al., 2021).

To further decipher the role of CK2 in cancer biology, we considered the opportunity of targeting allosteric sites of the protein. Allosteric compounds have the advantage to bind to less conserved pockets of the kinase. Conceptually, such compounds would be more selective chemical tools. Also, as therapeutics, they would generate less side effects due to off-target kinase inhibition (Lu et al., 2020). Regarding CK2, the CK2α/CK2β interface and the so-called αD pocket have both been targeted by small-molecule inhibitors (Figure S1) (Iegre et al., 2021). We previously reported a small-molecule that binds at the CK2α/CK2β interface and disrupts the holoenzyme in cells, inhibiting cell growth and migration (Kufareva et al., 2019). However, the affinity of the inhibitor was rather weak (K_D_ 30 μM) for further characterization. Bivalent CK2 inhibitors targeting simultaneously the ATP-binding site and the αD pocket have also been reported (Brear et al., 2016, De Fusco et al., 2017, Lindenblatt et al., 2022). The bivalent inhibitor CAM4066 (KD of 0.32 μM) reduced cell viability in HCT116, Jurkat and A549 cells with GI50 of 9, 6 and 20 μM, respectively (Brear et al., 2016). The bivalent inhibitor KN2 (K_I_ 6 nM for holoenzyme CK2) proved to be cytotoxic in HeLa cells (6 μM) but also displayed cytotoxicity in nontumor HEK293 cell line (16 μM). None of those bivalent compounds were used to explore CK2 function in cancer cells. Here, we report a highly selective CK2 bivalent inhibitor, AB668, that binds simultaneously the ATP site and the αD pocket of CK2α. AB668 displays an outstanding selectivity measured against a panel of 468 kinases, inhibits the activity of the holoenzyme CK2 with a K_i_ of 41 nM and induces apoptotic cell death on various cancer cell lines, while being non-cytotoxic for healthy cells. AB668 therefore represents a valuable chemical tool to explore CK2 mechanisms in cancer and to decipher the specific function of this particular pocket of the kinase. Here we compared the impact on cancer cells of AB668 and the two ATP-competitive CK2 inhibitors CX-4945 and SGC-CK2-1 on various cancer cells, using live cell imaging and transcriptomic analysis. Our results highlight that, by targeting both the ATP site and the αD pocket, AB668 discloses a distinct mode of action compared to canonical ATP-competitive inhibitors, suggesting that targeting the αD pocket of CK2 may represent a valuable pharmacological tool and therapeutic strategy in cancer.

## Results

### Binding mode, activity, and selectivity profile of AB668

During our efforts to optimize the CK2α/CK2β interface inhibitor that we previously reported (Kufareva et al., 2019, Figure S1), one of the chemical series led to the discovery of AB668, a compound that simultaneously binds at the ATP site and the αD allosteric pocket of CK2 (Figure 1a and Figure S2). A Thermal Shift Assay experiment was used to confirm direct binding of AB668 with CK2α. CK2α alone displayed a Tm of 43.6 ± 0.4°C. At saturating inhibitor concentration, AB668 induced a significant increase of the Tm value (ΔTm = 5.2 ± 0.4°C). By comparison, the ATP-competitive inhibitors CX-4945 and SGC-CK2-1 induced a substantial increase of the Tm value (ΔTm = 13.8 ± 0.4°C, ΔTm = 11.0 ± 0.3°C, respectively) (Figure S3). The lower magnitude of the change in Tm induced by AB668 may be related to its different binding site and to its lower affinity, as described below. The 3D X-ray structure of AB668 bound to CK2α was resolved using crystallization conditions reported for the X-ray structure of the CK2α-SGC-CK2-1 complex (Wells et al., 2021). The indole moiety of the bivalent inhibitor AB668 binds in the ATP site and interacts with the side chain of Lys68 through a hydrogen bond mediated by a water molecule, and the carbonyl group directly interacts with Lys68 through a weak hydrogen bond (Figure 1b). The aromatic moiety of indole is sandwiched between Val53, Val66, Ile95, Phe113, Met163 and Ile174. On the other side of the bivalent inhibitor, the substituted phenyl binds in the highly hydrophobic αD-pocket of CK2 (Tyr125, Leu128, Ile133, Met137, Tyr136, Ile140, Pro159, Val162, Ile164, Met221 and Met225). Regarding the linker part between the indole moiety and the substituted phenyl, the sulfonamide group interacts with the peptide chain of Ile164, while the nitrogen of piperidine interacts with Asn118 side chain. As illustrated in Figure 1c, the αD-helix position is shifted together with the side chains of residues Phe121 and Tyr125 upon AB668 binding and remains flexible in the crystal, with large B-factors values observed from residues Asn118 to Thr127. Modification of the conformation of the β4-β5 loop is observed in one monomer but not in the second one, suggesting that the loop conformation is highly dynamic, which is corroborated by large B-factors values for the loop.

**Figure 1.**
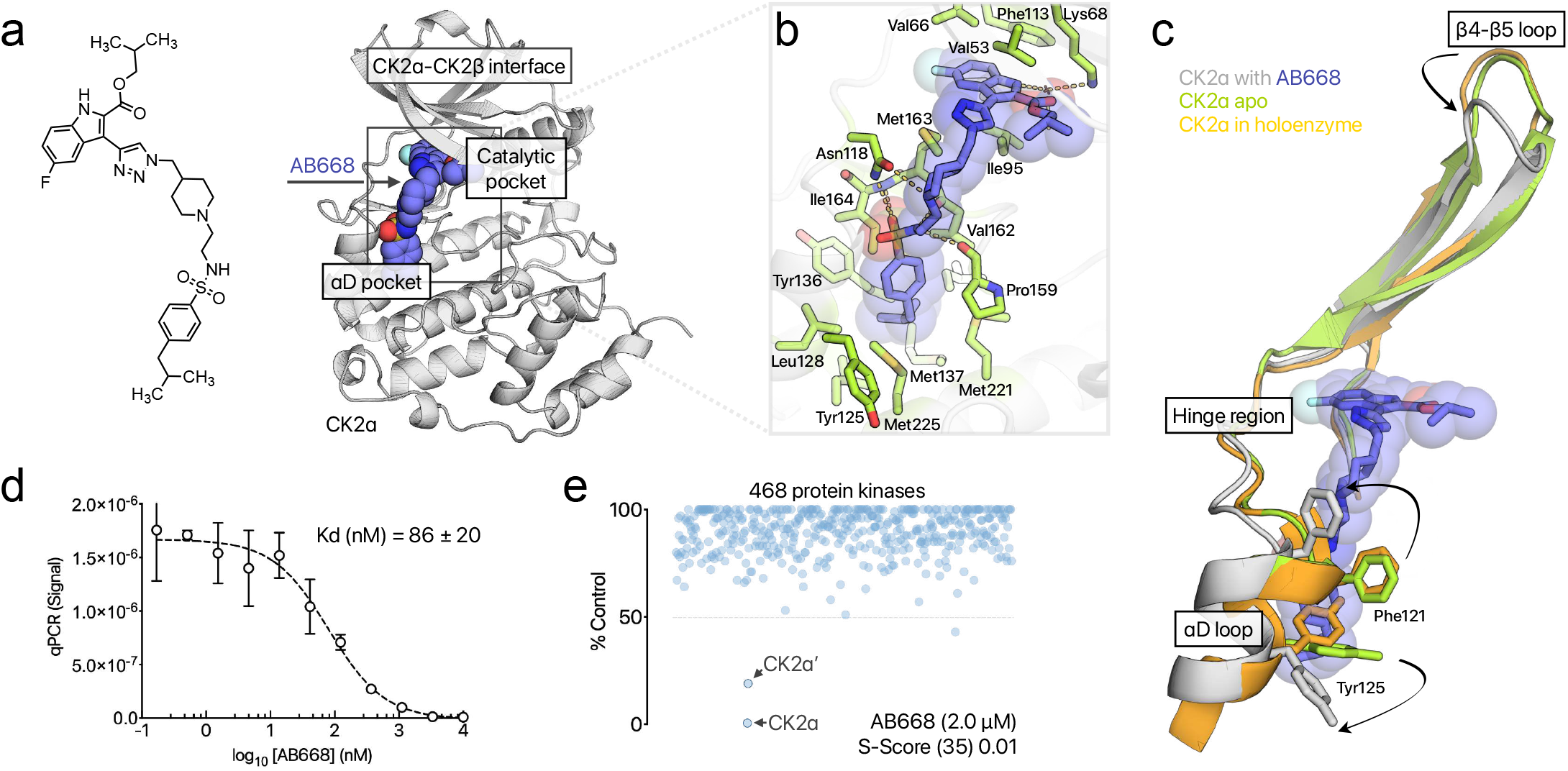
Binding mode, affinity for CK2α and kinase selectivity profile of AB668. **a**) Chemical structure of AB668 and CK2α/AB668 complex crystal 3D structure showing the binding site of AB668: the inhibitor binds both the ATP pocket and the αD pocket of CK2α. **b**) Stick representation of AB668 bound to CK2α: side chains in interaction with the inhibitor are shown, and hydrogen bonds are displayed. **c**) Conformational rearrangement of CK2α upon AB668 binding: the helix is shifted to allow the binding of AB668 in the αD pocket. The structure of CK2α bound to AB668 is superimposed to the structure of the holoenzyme (pdb code 1JWH), and to the apo structure of CK2α (pdb code 3QAO) **d**) Binding affinity of AB668 as determined by the KINOME*scan*™ profiling service (Eurofins). **e**) Selectivity profile of AB668, profiled against 468 kinases, using the screening platform from Eurofins DiscoverX. AB668 concentration was 2 μM (25 times its K_d_ value).

The affinity of AB668 for CK2α was 86 ± 20 nM, as determined by the KINOME*scan*™ profiling service (Eurofins) (Figure 1d). AB668 also inhibited the CK2 holoenzyme with an inhibitory constant Ki of 41 nM, as determined using a canonical radiometric assay with a CK2β-dependent peptide substrate (Figure S4a). We also showed that AB668 inhibits the phosphorylation of a CK2 protein substrate. The phosphorylation of SIX1, a transcription factor that is specifically phosphorylated by the CK2 holoenzyme (Wu et al., 2016) was inhibited by about 90% in the presence of 0.5 µM AB668 (Figure S4b).

The selectivity of AB668 was profiled against 468 kinases, using an active site-directed competition binding assay and IC_50_ validation with the screening platform from Eurofins DiscoverX. For this, AB668 was tested at a concentration of 2 μM (25 times its K_d_ value). As shown in Figure 1e, besides CK2α and CK2α’, only one kinase (RPS6KA5) displayed a percent inhibition larger than 50%. A value of 0.01 was obtained for the selectivity score (S10(2 μM)) (Bosc et al., 2017), showing that, by targeting the ligandable αD pocket, AB668 displays an outstanding selectivity against a large kinase panel.

### Cellular activity of AB668 in cancerous cells

We next evaluated whether this allosteric inhibitor was capable of engaging and inhibiting CK2 in living cells. As we previously reported that the CK2 subunits are overexpressed at the protein level in renal carcinoma compared to normal renal tissues (Roelants et al., 2015), we tested the effects of AB668, CX-4945 and SGC-CK2-1 in clear cell renal cell carcinoma (ccRCC) 786-O cells. Like staurosporine, AB668 and CX-4945 induced caspase-3 activation in 786-O cells, as shown after 72 h treatment using a quantitative fluorometric assay (Figure 2a). In contrast, and as previously reported (Wells et al., 2021), SGC-CK2-1 did not activate caspase-3 (Figure 2a). Caspase-3 activation induced by AB668 was confirmed by Western blot experiments (Figure 2b), showing the cleavage of PARP, a known target of caspase-3 (Duncan et al., 2011; Boulares et al., 1999). Similarly, AB668 reduced the expression of survivin, a member of the Inhibitor of Apoptosis Protein family that inhibits caspases and blocks cell death (Figure 2b) (Barrett et al., 2011). These observations encouraged us to further characterize the potential functional impact of AB668 and the ATP-competitive inhibitors in living cancer cells.

**Figure 2.**
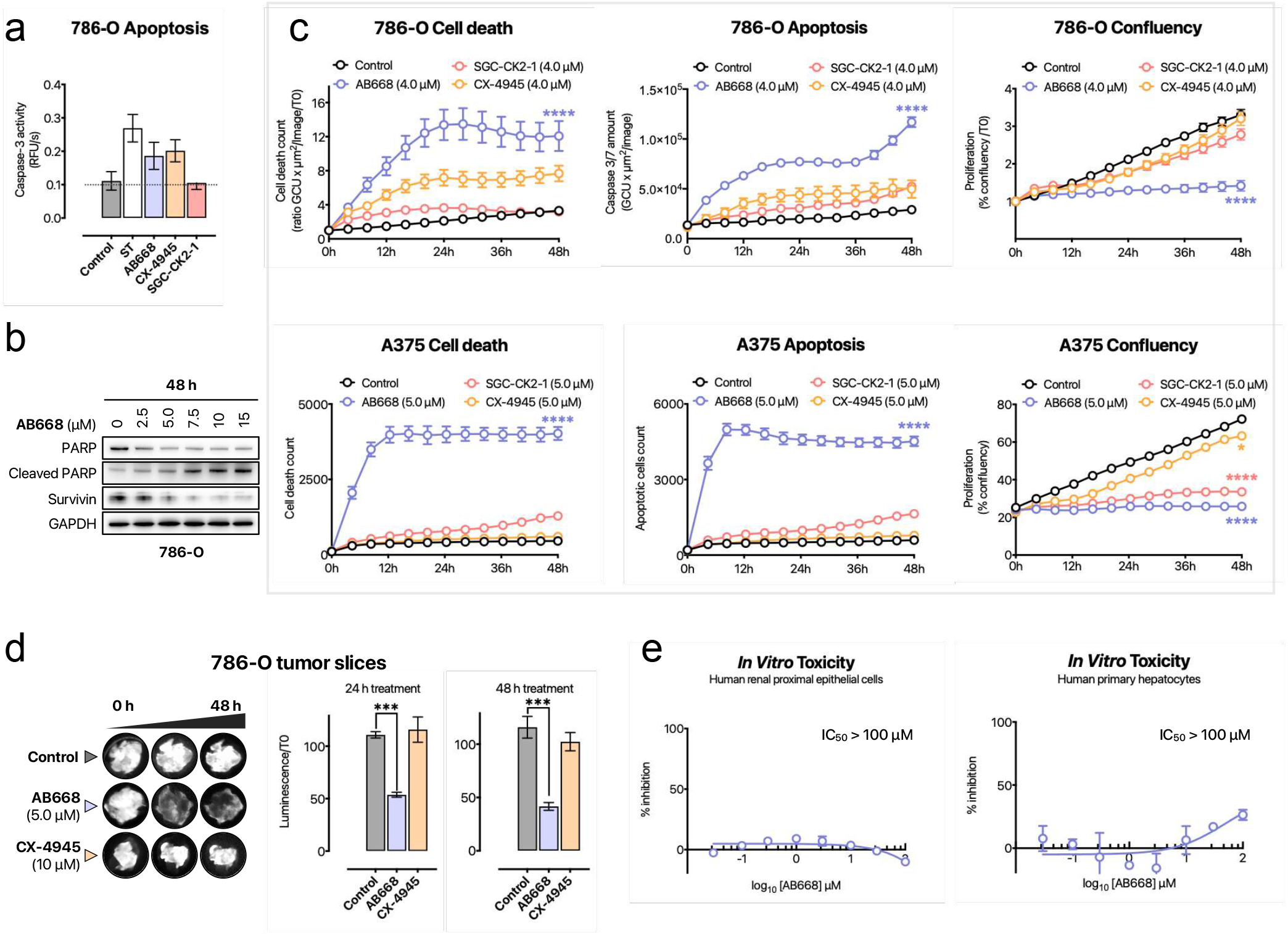
Cellular activity of AB668 and comparison to CX-4945 and SGC-CK2-1. **a**) Quantitative fluorometric assay showing caspase-3 activation in 786-O cells treated with staurosporine, AB668, SGC-CK2-1 or CX-4945. Assays were performed after 72 h treatment with the compounds at 20 µM. **b**) Western blot experiments on 786-O cells treated with AB668 (2.5, 5, 7.5, 10 and 15 µM) for 48h showing the cleavage of PARP as well as the expression level of survivin. **c**) Live cell imaging showing proliferation arrest, cell death and apoptosis in 786-O and A375 cells treated with AB668, CX-4945 or SGC-CK2-1 (4 µM) for 48h. **d**) Effect of AB668 and CX-4945 on e*x vivo* culture of intact tumor slices of clear cell renal carcinoma. Tumors were extracted from renal carcinoma xenografted mice that were directly processed into 300 μm slices and treated for 48 h as described in Methods. Cell viability was evaluated by luciferin measurement of treated tumor-slice cultures as described in Methods. **e**) Cell viability of primary hepatocytes and RPTEC (Renal Proximal Tubule Epithelial Cells) treated with AB668.

For this, Incucyte Live cell analysis was used to analyze the effect of the various CK2 inhibitors in 786-O cells (Figure 2c). We also performed the same experiments for a melanoma cell line (A375), for which few information is available regarding the effect of CK2 inhibition (Parker et al., 2014; Zhou et al., 2016). As illustrated in Figure 2c, AB668 induced significant cell proliferation arrest associated with cell death and apoptosis in both cancer cell lines. By comparison, CX-4945 had only a moderate effect on 786-O cell growth and was a poor apoptosis inducer. Similarly, A375 cells were almost completely insensitive to CX-4945 and SGC-CK2-1 showing that these inhibitors had no impact on the survival of both cell lines (Figure 2c). CK2 has been shown to be involved in apoptotic pathway protecting substrates from caspase-3-mediated proteolysis (Duncan et al., 2011; Turowec et al., 2013), and CX-4945 was previously reported to induce apoptotic cell death in cancer cells lines such as PC3 prostatic adenocarcinoma (Pierre et al., 2011), B-ALL, T-ALL (Richter et al., 2019; Buontempo et al., 2014), H1299, Calu-1 and H358 (So 2015). Our data suggest that the selective CK2 inhibition by SGC-CK2-1 is not sufficient to induce caspase-3-mediated apoptosis and that apoptotic cell death induced by CX-4945 may be related to its activity on other kinases.

We then evaluated the efficacy of AB668 on an *ex-vivo* culture model of renal carcinoma that we previously used to study individual responses to targeted therapies (Roelants et al., 2018; Roelants et al., 2020). As shown in Figure 2d, 10 µM CX-4945 had no effect on tumor slices of renal carcinoma, while 5 µM AB668 significantly reduced cell viability after 24h of treatment, showing its higher efficacy in this drug sensitivity prediction model. This study suggests that AB668-mediated CK2 inhibition could prove a viable therapeutic strategy in renal carcinoma and highlights that treatment of tumor tissue slice culture best predicts CK2 dependency.

Importantly, no cytotoxicity of AB668 was observed in normal human cell lines such as RPTEC (Renal Proximal Tubule Epithelial Cells) and primary hepatocytes (Figure 2e).

### Target engagement of AB668 in cancer and healthy cells

We next evaluated the effects of AB668 in 786-O cells on CK2-mediated downstream phosphorylation events by western blot analysis. Notably, CK2 is known to phosphorylate AKT at Ser129 (Di Maira et al., 2005; Zanin et al., 2012). As expected, AB668 decreased, in a dose-dependent manner, the phosphorylation of S129AKT (Figure 3a). AB668 also induced a dose-dependent inhibition of the activated forms of mTOR and STAT3, two proteins that regulate proliferation and apoptosis in cancer cells (Sabatini et al., 2006; Yuan et al., 2004). It was reported that inhibition of CK2 hinders STAT3 signaling and decreases aggressive phenotypes in multiple cancer types (Gray et al., 2014; Aparicio-Siegmund S. et al., 2014). AB668 activated p38MAPK, a tumor suppressor well known for its role in transducing stress signals from the environment (Figure 3a). Interestingly, mouse modeling studies showed that mTOR activation in combination with inactivation of the p38MAPK initiates renal cell carcinoma (Wu et al., 2021).

**Figure 3.**
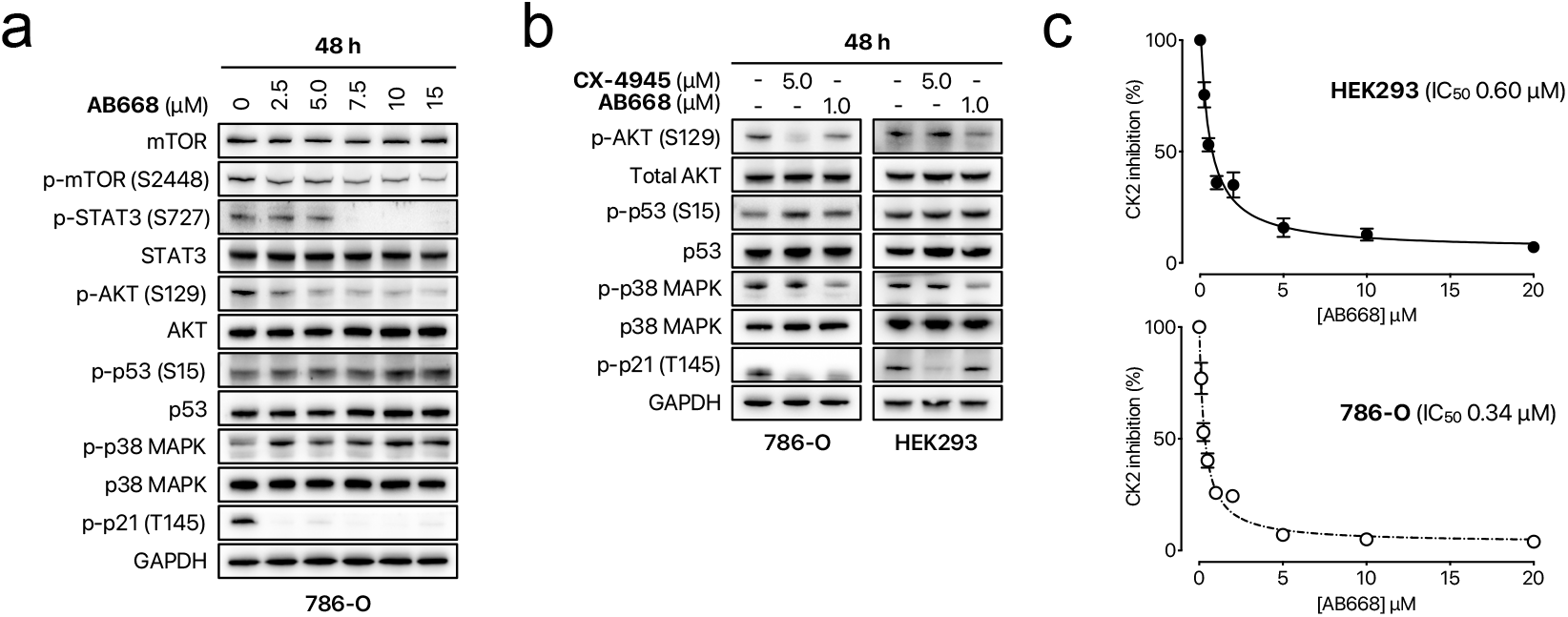
Target engagement of AB668 in 786-O cells and HEK293 cells. **a**) Western blot analysis in response to AB668 in 786-O cells. The inhibition of phosphorylated mTOR (S2448), STAT3 (S727), p53 (S15), p38 MAPK (T180/Y182) and AKT (S129) and p21 (T145) was analyzed after 48 h of inhibitor AB668 treatment. **b**) The inhibition of phosphorylated AKT (S129), p53 (S15), p38 MAPK (T180/Y182) and p21 (T145) was analyzed after 48 h of AB668 (1 µM) or CX-4945 (5 µM) treatment in 786-O cancer cells or HEK293 cells. Glyceraldehyde-3-phosphate dehydrogenase (GAPDH) was used as a loading control. **c**) CK2 activity measured in cell extracts of 786-O cells (upper panel) or HEK293 cells (lower panel) after treatment with AB668. Inhibition constants IC_50_ are 0.34 ± 0.07 µM for 786-O cells and 0.60 ± 0.11 µM for HEK293 cells.

Although p53 is rarely mutated in ccRCC, its overexpression has been linked to poor prognosis (Noon et al., 2010; Diesing et al., 2021). Similarly, it has been reported that irradiation of ccRCC cell lines including 786-O cells, induced p53 phosphorylation without detectable activation indicating a functional inhibition in ccRCC (Diesing et al., 2021). As illustrated in Figure 3a, AB668 led to a weak increase of p53 phosphorylation only at high concentrations, ruling out its implication in the regulation of 786-O cell viability.

Previous work has demonstrated the anti-apoptotic role of the cyclin-dependent kinase inhibitor p21 in RCC as a potential mechanism for their drug resistance (Weiss et al., 2003). p21 binds to and inhibits the activity of proteins involved in apoptosis, including pro-Caspase-3 (Karimian et al., 2016; Abbas, et al., 2009). Phosphorylation of p21 at Thr145 by AKT1 induces its cytoplasmic accumulation (Rossig et al., 2001; Zhou et al., 2001) and propels its anti-apoptotic functions (Karimian et al., 2016; Abbas, et al., 2009). Interestingly, Sorafenib, as one of the few available effective therapeutic options for metastatic RCC, was shown to attenuate the anti-apoptotic role of p21 in kidney cancer cells (Inoue et al., 2011). We have previously shown that CX-4945 inhibits the AKT1-dependent phosphorylation of p21 in 786-O cells (Roelants et al., 2015). As shown in Figure 3a, this phosphorylation was also strongly downregulated by low concentrations of AB668 in accordance with its inhibitory effect on AKT phosphorylation in 786-O cells. Finally, we have also evaluated the effects of AB668 in the triple-negative breast cancer cell line MDA-MB231 by western blot analysis. In comparison with 786-O cells, AB668 led to similar effects on CK2-mediated downstream phosphorylation events (Figure S5).

Motivated by the striking difference in sensitivity to AB668 between cancer versus normal cells, we evaluated its target engagement by assaying the CK2 activity in extracts of 786-O cancer cells and human embryonic kidney HEK293 cells after treatment with increasing concentrations of AB668. Inhibition constants IC_50_ were 0.34 ± 0.07 µM and 0.60 ± 0.11 µM for 786-O cells and HEK293 cells, respectively (Figure 3c). These results show that AB668 inhibited CK2 activity with a similar potency in both cell types although it did not induce cytotoxicity in normal cells (Figure 2e and Figure S5b). The comparison of downstream phosphorylation events mediated by CK2 in 786-O and HEK293 cells in response to CX-4945 or AB668 indicates that p21 phosphorylation was impaired in 768-O cells but unaffected in HEK293 cells in response to AB668 (Figure 3b).

### Deregulations of the transcriptome in response to AB668 or CX-4945

To assess the impact of AB668 and CX-4945 at the transcriptomic level, molecular profiling of 786-O cells treated with these CK2 inhibitors was performed by BRB-seq, followed by differential gene expression analysis (Figure 4a – left and center plot). Only 48 genes were significantly differentially expressed (Benjamini-Hochberg corrected p-value < 0.05 and |log_2_ (Fold change)| > 0.5) in response to AB668 when compared to cells treated with DMSO. In contrast, 102 genes were deregulated by CX-4945 compared to DMSO. This might be related to the lower specificity of this CK2 inhibitor. Moreover, 341 genes were significantly differentially expressed in cells exposed either to AB668 or CX-4945, revealing strong differences in transcriptome perturbations associated to these two CK2 inhibitors (Figure 4a – right plot). Interestingly, SERPINE1, also known as Plasminogen Activator Inhibitor type-1 (PAI-1) a protein with a growth and migration stimulatory functions and an anti-apoptotic activity activity (Planus et al., 1997; Kubala and DeClerck, 2019) was found significantly down-regulated by AB668, when compared to CX-4945 or DMSO. In ccRCC tumor tissues, the expression level of PAI-1 is higher than in normal tissues and has been proven to be a reliable biological and prognostic marker associated with poor prognosis (Sui et al., 2021). Metabolic enzymes such as PSAT1, the phosphoserine aminotransferase (PSAT1), UDP-N-acetylglucosamine pyrophosphorylase 1 (UAP1) the UDP-N-acetylglucosamine pyrophosphorylase 1 and methylenetetrahydrofolate deshydrogenase/cyclohydrolase (MTHFD2), the methyleneterahydrofolate deshydrogenase/cyclohydrolase that are highly expressed in a wide range of tumors and associated with poor prognosis in tumor progression (Itkonen, Engedal et al., 2015; Puttamallesh et al., 2020; Reina-Campos et al., 2020; Huang et al., 2021; Zhu et al., 2022) were significantly down regulated by AB668 when compared to CX-4945.

**Figure 4.**
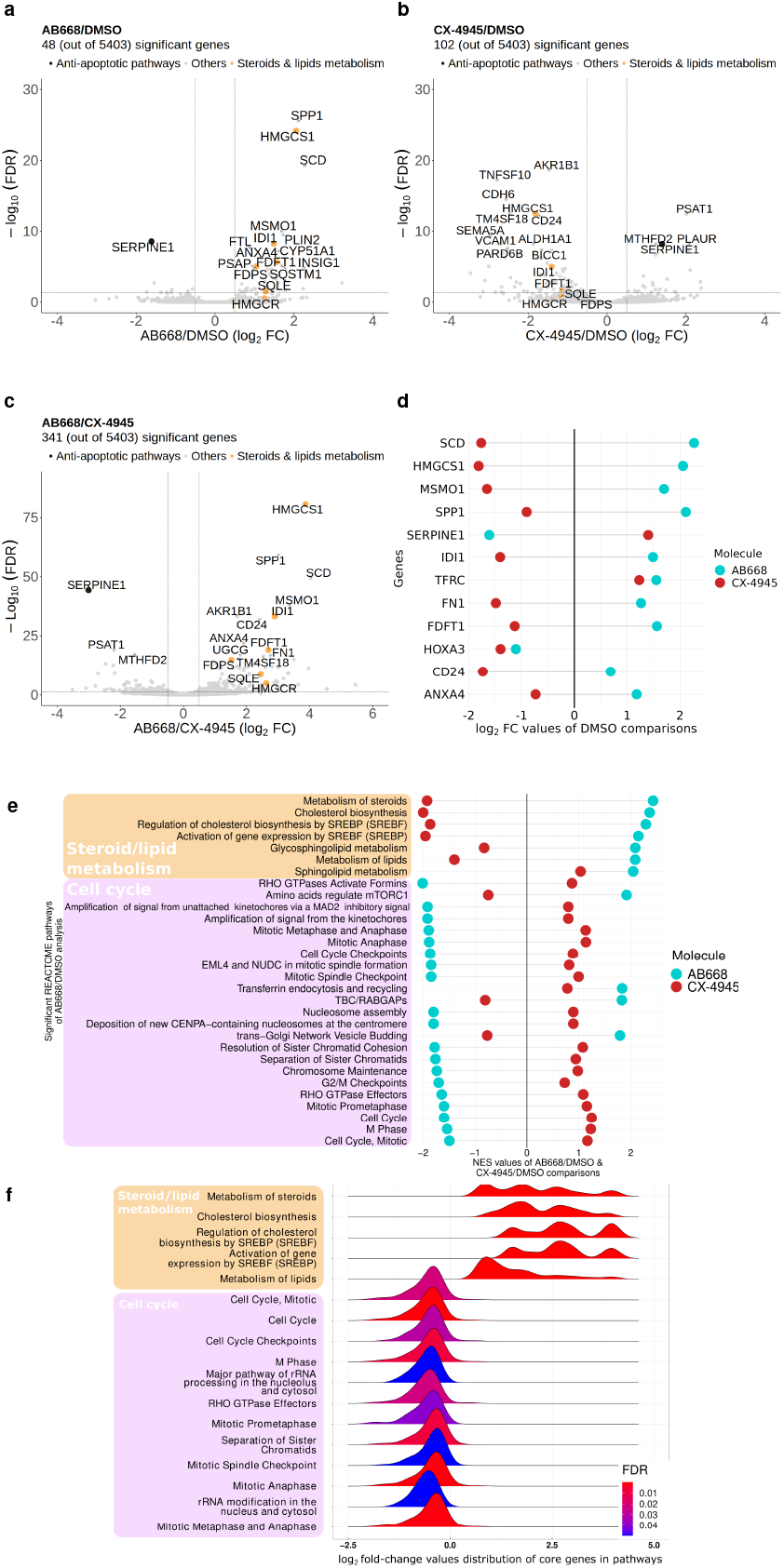
Deregulations of the transcriptome in response to AB668 or CX-4945. **a-c)** Volcano plots based upon differential gene expression analysis of the transcriptomes of 786-O cells exposed to the different CK2 inhibitors: **a)** AB668 *vs* DMSO **b)** CX-4945 vs DMSO and **c)** AB668 *vs* CX-4945. **d)** Differences of Log2(Fold Changes) of gene expression values for AB668 and CX-4945 treated cells compared to DMSO treated cells. **e)** Normalized Enrichment Scores (NES), calculated by the Gene Set Enrichment Analysis (GSEA) method, for significantly deregulated pathways obtained from AB668 *vs* DMSO analysis in comparison to the NES scores generated from CX-4945 *vs* DMSO analysis. **f)** Histograms of Log2(Fold Changes) of gene expression values for AB668 *vs* CX-4945 comparison for significantly deregulated pathways found by GSEA analysis, using REACTOME pathway database.

In addition, we specifically identified, among the genes differentially expressed compared to the DMSO control, those whose expression was deregulated in an opposite way between AB668 and CX-4549 (Figure 4b). For example, SERPINE1 is down-regulated by CX-4945 but up-regulated by AB668 compared to DMSO-exposed cells. Interestingly, genes involved in fatty acid, cholesterol and steroid metabolisms like, Farnesyl-diphosphate farnesyltransferase 1 (FDFT1), 3-hydroxy-3-methylglutaryl-CoA synthase 1 (HMGCS1) and Isopentenyl-diphosphate delta isomerase 1 (IDI1) were all up-regulated by AB668 while down-regulated by CX-4945. Other fatty acid metabolism related genes 3-Hydroxy-3-Methylglutaryl-CoA Reductase (HMGCR) and Squalene Epoxidase (SQLE) were also up-regulated in response to AB668 compared to CX-4945 or DMSO (Figure 4a). This can be related to the recently pointed out role of CK2 in lipid homeostasis and its deregulation in cancer cells (Guerra et al., 2020). In particular, HIF expression drives lipid deposition in ccRCC *via* the repression of fatty acid metabolism (Du et al., 2017). Importantly, it was previously shown that low expression of FDPS, FDFT1, HMGCS1, HMGCR and IDI1 genes and high expression of SQLE were associated with patients with high-risk of ccRCC (Qi et al., 2021). Thus, by up-regulating 5 out of 6 genes in this prognostic signature, AB668 could represent a therapeutic option to counteract ccRCC tumor progression.

An identification of the biological pathways over-represented in genes deregulated in expression, in response to either AB668 or CX-4945 compared to DMSO, was then carried out using the Gene Set Enrichment Analysis (GSEA) method and the Reactome database. Among the 17 significantly altered pathways, 5 were related to the metabolism of steroids and lipids and 12 were connected to cell cyle and mitotic processes (Figure 4d). While pathways related to steroids and lipids metabolism were up-regulated in response to ABB668, cell cycle and mitotic processes-based pathways were down-regulated by AB668. This down-regulation of cell cycle is consistent with the strong inhibition of cell proliferation observed in cell lines treated with AB668. Furthermore, among the pathways identified as significantly deregulated in response to AB668 compared to DMSO, we found that all but two (transferin endocytosis and recycling and sphingolipid metabolism) have opposite deregulations in response to CX-4945 (Figure 4c), illustrating the strongly different effects of these CK2 inhibitors on these cellular processes.

In summary, our transcriptomic analysis clearly demonstrates that the inhibition of CK2 by the dual inhibitor AB668 or by a pure ATP-competitive CX-4945 inhibitor that reached clinical trials differentially affect biological pathways in 786-O cancer cells. More specifically, AB668 induces a deep alteration of antiapoptotic pathways, cell cycle and mitotic processes, as well as steroids and lipids metabolism, which all are involved in renal cancer tumorigenicity.

## Discussion

The key role of CK2-dependent pathways in cancer has motivated the development of CK2 inhibitors. While most of these inhibitors act by an orthosteric mechanism, meaning that they bind to the highly conserved ATP binding-pocket of the kinase (Cozza et al., 2012), small molecules that act outside the ATP site have been also described (Iegre et al., 2021). Here, we have disclosed a new bivalent CK2 inhibitor that binds at both the ATP site and the allosteric αD pocket, a feature that accounts for its high selectivity profile in the human kinome. The presence of this ligandable allosteric pocket on CK2 was previously revealed during a crystallographic fragment screening campaign (Brear et al., 2016), in agreement with the high mobility of the αD helix in CK2. The flexibility of this αD helix was experimentally observed in various crystallographic structures of the kinase and was also reported in metadynamic studies of CK2 structure (Gouron et al., 2014). Two bivalent CK2 inhibitors targeting this pocket, in addition to the ATP site, have been previously published (Brear et al., 2016; Lindenblatt et al., 2022). While their impact on cell cytotoxicity has been reported, the effect of such inhibitors on cellular CK2 functions has not been explored.

Here, we have compared the effects of the bivalent CK2 inhibitor AB668 to CX-4945 and SGC-CK2-1 using caspase activation assay, live-cell imaging and transcriptomic analysis. Our data highlight that AB668 has a distinct mechanism of action regarding its anti-cancer activity. Treatment with AB668 strongly impacted cancer cell viability resulting in apoptotic cell death, while sparing healthy cells. A long-held view has suggested that malignant cells are dependent upon sustained CK2 signaling for survival thereby exhibiting an exquisite sensitivity to CK2 inhibition (Ruzzene et al, 2010; Roffey et al, 2021; Firnau et al, 2022). Conversely, non-cancer cells are more resistant to induction of cell death on downregulation of CK2 activity, which is the expected basis for safely using a pharmacological chemical inhibitor. By contrast, although CX-4945 or SGC-CK2-1 reduced cell proliferation, both compounds were inefficient to induce cell death. These observations suggest that targeting the allosteric CK2 αD pocket has a distinct cellular impact as compared to targeting only the CK2 ATP binding site. These results also indicate that CK2 is a key player in cancer biology and that targeting CK2 is a valuable strategy (Salvi et al., 2021).

Transcriptomic analysis further supports this hypothesis, as we could highlight striking differences in biological pathways induced by cell treatment with either AB668 or the ATP-competitive inhibitor CX-4945. AB668 acts by an unconventional mechanism and induces strong apoptotic cell death. Thus, this strong pharmacological effect observed in various functional assays indicates that AB668 is an important novel investigational probe for exploiting apoptotic vulnerabilities in cancer as well as a promising lead for the next generation of drug-like CK2 inhibitors with improved potency and optimal drug properties. Future experiments will need to define further the mechanisms by which AB668, by occupying the allosteric αD pocket, induces apoptotic cell death in cancer cells, while sparing healthy cells. Taken together, our results strongly suggest that CK2 inhibition using small molecules that target binding sites outside the ATP pocket could be a valuable strategy in treating various aggressive cancers and disease-relevant contexts.

## Methods

### Recombinant proteins for enzymatic measurements

Both human recombinant CK2α subunit and chicken recombinant MBP (maltose-binding protein)-CK2β were expressed in *Escherichia coli* and purified as previously reported (Hériché et al., 1997; Chantalat et al., 1999). Proteins were quantified using a Bradford assay and the quality of the purification was asserted by SDS-PAGE analysis.

### CK2 activity assays in vitro

Radiometric kinase assays were performed as previously reported (Kufareva et al., 2019). Briefly, in a final volume of 20 μL at 4°C, 3.0 μL of CK2α protein (36 ng) was incubated in the reaction mixture (20 mM Tris-HCl, pH 7.5, 150 mM NaCl, 1.0 mM DTT) with 1.0 mM of the synthetic substrate peptide, 20 mM of MgCl_2_, 1.0 μCi of [^32^P]-ATP and 2.0 μL of different concentrations of the inhibitor, diluted in Tris-HCl-glycerol, 0.05% Tween 20. Final ATP concentration was 10 μM when not stated otherwise. The kinase reactions were performed under linear kinetic conditions for 5 min at room temperature followed by quenching with the addition of 60 μL of 4% TCA. ^32^P incorporation in peptide substrate was determined by spotting the supernatant onto phospho-cellulose paper disks (Whatman P81, 4 cm^2^). The disks were washed three times in cold 0.5% phosphoric acid, 5 min on a rocking platform per wash, dried and finally the radioactivity was measured. Percentage inhibition was calculated relative to a DMSO control and all measurements were performed in duplicate. A canonical CK2 peptide substrate (Seq. RRREDEESDDE) phosphorylated equally by CK2α_2_β_2_ (CK2β-independent) and a 22-residue long *N*-terminal fragment of the eukaryotic translation initiation factor 2 (eIF2) (.MSGDEMIFDPTMSKKKKKKKKP), exclusively phosphorylated by CK2α_2_β_2_ (CK2β-dependent) were used for the radiometric kinase assays. Phosphorylation assay using GST-SIX1 (3.7 μg) were performed in the same buffer. Final concentration of ATP was 100 μM. Samples were analyzed by SDS-PAGE and subjected to autoradiography. Phosphoproteins were quantified by densitometry scanning using ImageJ (National Institutes of Health software v1.52).

### X-ray crystallography

Recombinant protein for X-ray studies was produced as published in Wells et al. (2021). Protein concentration was 9 mg/ml. Crystals of human CK2α were grown at 20 °C using the hanging-drop vapor-diffusion method with a reservoir solution containing 33% polyethylene glycol methyl ether 5000, 0.2 M ammonium sulfate, 0.1 MES pH 6.5. The drops contained 1µl of the reservoir solution and 1 µl of the protein. The crystals were soaked by adding 0.2 µl of ligand solution at 100 mM in DMSO. The crystals were cryo-protected with reservoir solution supplemented with 20% glycerol and then flash-cooled in liquid nitrogen. X-ray diffraction data were collected at the ESRF Synchrotron in Grenoble, France, on beamline IB30B. Data were integrated and processed using XDS (Kabsch et al., 2010). The crystals belong to the space group P43212 with two monomers in the asymmetric unit. The structures were solved by molecular replacement using PDB entry 6Z84 as the search model. Bound ligands were manually identified and fitted into Fo–Fc electron density using Coot (Emsley & Cowtan, 2004). Files CIF format for ligand were generated using Grade Server (http://grade.globalphasing.org/cgi-bin/grade/server.cgi). The structure was refined by rounds of rebuilding in Coot and refinement using Phenix (Adams et al., 2010). Data collection and refinement statistics for crystal structure is presented in Table S1.

### Thermal Shift Assay

The thermal shift assay was performed on a LightCycler 480 Real-Time PCR System (Roche) in 96-well white plates (Armadillo plate, Thermo Scientific) using an integration time of 120ms. Each well contained 10 μL of 5µg CK2α (purified as described by Hériché et al., 1997; Chantalat et al., 1999) and 2.5× SYPRO Orange (Life Technologies) in PBS-0.9% glycerol, with ligands added to a final concentration of 0.1µM to 500µM in 5% (v/v) DMSO. All assays were carried out in triplicate. Each plate was sealed with an optically clear foil and centrifuged for 1 min at 300 rpm before performing the assay. The plates were heated from 20 to 80°C at a heating rate 0.01°C/s. The fluorescence intensity was measured with λex = 483 nm and λem = 568 nm. The melting temperature (Tm) was determined using the TSA-CRAFT software that enables automatic analysis of TSA data exported from the Roche Lightcycler 480 software (Lee et al., 2019).

### Kinase screening

Profiling of 468 recombinant protein kinases was performed by Eurofins Discovery (KINOMEscan™ Profiling Service, San Diego, USA) at 10 μM of ATP in the presence of 2 μM AB668. Inhibition, expressed as the percent of activity, was calculated from the residual activity measured in the presence of the inhibitor.

### Cell lines

All cell lines were purchased from American Type Culture Collection (ATCC) and grown on standard tissue culture plastic in a 5% CO_2_ humidified incubator at 37°C. 786-O were maintained in RPMI 1640 medium (Gibco), containing 10% of FBS, penicillin (24 U/mL), and streptomycin (25 μg/mL). A549, A375 and MDA-MB231 were cultured in DMEM + GlutaMAX medium (Gibco) supplemented with 10% of FBS. HEK293T were grown in EMEM supplemented with 10% of FBS and RPTEC were maintained in ProXup (Evercyte). MCF10A were cultured as described (Debnath et al., 2003).

### Caspase-3 assay

The Caspase-3 Fluorometric Assay Kit II from Biovision was used to determine caspase-3 activity in cells following manufacturer’s instructions. Briefly, cultured cells collected by scraping were lyzed and Bradford reagent was used to quantify proteins in the cell lysate. Fifty micrograms of proteins were added in a 96-well white plate. Reaction was started by adding the caspase-3 substrate, DEVD-AFC. Fluorescence measurements (excitation: 405 nm; emission: 520 nm) were made at 37°C in a FLUOstar OPTIMA microplate reader (BMG Labtech) every 5 min over an hour. Cells treated with 500 nM staurosporine, a potent apoptotis inducer, were used as positive controls.

### Cell death & proliferation

786-O (2 × 10^4^ cells per well), HEK293 (2 × 10^4^ cells per well), MCF10A (2 × 10^4^ cells per well), MDA-MB231 (2 × 10^4^ cells per well) and RPTEC (7 × 10^3^ cells per well) cells were seeded into 96-well flat-bottom cell culture plates. After 24 h, compounds dissolved in DMSO were diluted in the culture medium containing 0.5 μg/mL propidium iodide (PI, Sigma-Aldrich) and were added to the cell culture media such as the final DMSO concentrations is equal to 0.2% (v/v). Experiments were conducted at 37°C in a 5% CO2 atmosphere and the plates were tracked using an Essen IncuCyte Zoom live-cell microscopy instrument. For cell death, PI-stained red fluorescent cells images were captured every 3 h for the entire duration of the experiment and normalized to the DMSO standard control. For cell proliferation, the software incorporated into the IncuCyte Zoom was specifically calibrated to ensure accurate distinction of cells from the empty spaces. Cell proliferation was monitored by analyzing the occupied area (% confluence) of cell images over time.

### Real-time proliferation, cell death and apoptosis assay on A375 cells

A375 melanoma cells (2.5 × 10^4^ cells per well) were plated in 48-well plates. After 24 hours, cells were treated with various doses of AB668, CX4945, or SGC-CK2-1. Treatment medium contained 0.3 µg/mL propidium iodide (Sigma-Aldrich) and 2 µM CellEvent™ Caspase-3/7 Green Detection Reagent (ThermoFisher Scientific). Images were captured automatically every two hours for 48 hours using the IncuCyte™ S3 Live-Cell Analysis Instrument (Essen BioScience). Image analysis was performed using IncuCyte software. Data was plotted as mean ± SD using GraphPad Prism 9.

### Preparation of cell extracts

786-O (3 × 10^5^ cells per well), HEK 293 (2 × 10^5^ cells per well), MDA-MB231 (3 × 10^5^ cells per well), and RPTEC (3 × 10^5^ cells per well) cells were seeded into 6-well tissue and cultured for 24 h prior to the addition of inhibitors (as described above). After incubation, medium was removed, cells were washed with cold PBS and frozen at −80°C. Cells were lysed using a RIPA buffer (10 mM Tris-HCl, pH 7.4, 150 mM NaCl, 1% Triton X-100, 0.1% SDS, 0.5% DOC and 1.0 mM EDTA) with the addition of protease and phosphatase inhibitor cocktail (Sigma-Aldrich, P8340, P2850, P5726) at the recommended concentrations. Cell pellets were incubated in RIPA buffer on ice for 30 min, then centrifuged for 15 min at 4°C at 13000 rpm and the supernatants collected. Proteins were quantified using the Pierce BCA protein assay kit (Pierce, ThermoFisher Scientific).

### CK2 activity in cells

Cell homogenates were assayed for CK2 activity with radiometric assays as described above.

### Immunoblotting

Equal amounts of lysates (20-35 μg) were loaded onto a precast 4-12% gradient gel (Bio-Rad) and submitted to electrophoresis in NuPAGE buffer (150 V for 1.5 h). The gels were transferred onto PVDF membrane (100 V for 1 h). Membranes were blocked during 1 h at room temperature with saturation buffer (5% BSA in Tris-buffered saline with 0.1% Tween 20 (TBST) and then incubated with primary antibody diluted in saturation buffer for 2 h or overnight on a rocking platform shaker. Primary antibodies were GAPDH antibody (#AM4300) from Invitrogen, ThermoFisher, P-AKT-phospho-Ser129 (#AP3020a) from Interchim, AKT (#9272), PARP (#9542), mTOR (#2972), mTOR-phospho-Ser2448 (#2971), p38MAPK (#9212), p38MAPK-phospho-Thr180/Tyr182 (#9211), p53 (#9282), p53-phospho-Ser15 (#9286), STAT3 (#6139), STAT3-phospho-Ser727 (#9134) antibodies from Cell Signaling, survivin (#NB500201) antibody from Novus biologicals, p21 antibody (#sc-397) from Santa Cruz Biotechnologies, p21-phospho-Thr145(#ab-47300) antibody from Abcam. After washing three times with TBST, secondary antibody (peroxidase-conjugated affinity pure Goat anti-rabbit IgG (#111035003) or peroxidase-conjugated affinity pure goat anti-mouse IgG (#115035003) from Jackson Immuno Research) was added for 1 h followed by three more washes with TBST. Immobilon Forte Western HRP substrate (Millipore) was added and detection was achieved by using Fusion FX acquisition system (Vilbert). Anti-GAPDH was used as loading control and images were analyzed and band intensities were quantified using ImageJ (National Institutes of Health software v1.52).

### *In vivo* orthotopic tumor xenograft models

All animal studies were approved by the institutional guidelines and those formulated by the European Community for the Use of Experimental Animals. Six week-old BALB/c Female nude mice (Charles River Laboratories) with a mean body weight of 18-20 g were used to establish orthotopic xenograft tumor models. The mice were housed and fed under specific pathogen-free conditions. To produce tumors, renal cancer cells 786-O-luc were harvested from subconfluent cultures by a brief exposure to 0.25 % trypsin-EDTA. Trypsinization was stopped with medium containing 10 % FBS, and the cells were washed once in serum-free medium and resuspended in 500 μl PBS. Renal orthotopic implantation was carried out by injection of 3 × 10^6^ 786-O luc cells into the right kidney of athymic nude mice. Mice were weighed once a week to monitor their health and tumor growth was measured by imaging luminescence of 786-O-luc cells (IVIS).

### Fresh tissue sectioning

A Vibratome VT1200 (Leica Microsystems) was used to cut thin (300 μm) slices from fresh tissue. Samples were soaked in ice-cold sterile balanced salt solution (HBSS), orientated, mounted, and immobilized using cyanoacrylate glue. Slicing speed was optimized according to tissue density and type; in general, slower slicing speed was used on the softer tissues and vice versa (0.08-0.12 mm/s neoplastic tissue; 0.01-0.08 mm/s normal tissue). Vibration amplitude was set at 2.95-3.0 mm.

### Organotypic tissue cultures

Tissue slices were cultured on organotypic inserts for 48 h (one slice per insert; Millipore). Organotypic inserts are Teflon membranes with 0.4 μm pores that allow preservation of 3D tissue structure in culture. Tissue culture was performed at 37°C in a 5 % CO_2_ humidified incubator using 1 ml of DMEM media supplemented with 20 % FBS (GIBCO), 100 U/ml penicillin (Invitrogen) and place in a rotor agitator to allow gas and fluids exchanges with the medium. The tissue slices were incubated with the CX-4945 (10 µM) or AB668 (5 µM) until 68 hours. Tumor slice viability was monitored at indicated time points, by luminescence imaging of 786-O-luc cells after luciferin addition (IVIS) as previously described (Roelants et al., 2018; Roelants et al., 2020).

### BRB-seq library preparation and sequencing

786-O cells (150, 000) were treated for 24h with DMSO, 5 µM of CX4945 or AB668. Total RNA was extracted from treated cells using the MirVana PARIS kit (Thermofisher). The 3′ Bulk RNA Barcoding and sequencing (BRB-seq) experiments were performed at the Research Institute for Environmental and Occupational Health (Irset, Rennes, France) according to the published protocol (Alpern et al., 2019). Briefly, the reverse transcription and the template switching reactions were performed using 4 µl total RNA at 2.5 ng/µl. RNA were first mixed with 1 µl barcoded oligo-dT (10 µM BU3 primers, Microsynth), 1 μL dNTP (0.2 mM) in a PCR plate, incubated at 65 °C for 5 min and then put on ice. The first-strand synthesis reactions were performed in 10 µl total volume with 5 µl of RT Buffer and 0.125 µl of Maxima H minus Reverse Transcriptase (Thermofisher Scientific, #EP0753) and 1 µl of 10 μM template switch oligo (TSO, IDT). The plates were then incubated at 42°C for 90 min and then put on ice.

After Reverse Transcription (RT), decorated cDNA from multiple samples were pooled together and purified using the DNA Clean & concentrator-5 Kit (Zymo research, #D4014). After elution with 20 µl of nuclease free water, the samples were incubated with 1 µl Exonuclease I (NEB, #M0293) and 2 µl of 10X reaction buffer at 37°C for 30 min, followed by enzyme inactivation at 80°C for 20 min.

Double-strand (ds) cDNAs were generated by PCR amplification in 50 µl-total reaction volume using the Advantage 2 PCR Enzyme System (Clontech, #639206). PCR reaction was performed using 20 µl cDNA from previous step, 5 µl of 10X Advantage 2 PCR buffer, 1 µl of dNTPs 50X, 1 µl of 10 µM LA-oligo (Microsynt), 1 µl of Advantage 2 Polymerase and 22 µl of nuclease-free water following the program (95°C-1 min; 11 cycles: 95°C-15 s, 65°C-30 s, 68°C-6 min; 72°C-10 min). Full-length double-stranded cDNA was purified with 30 µl of AMPure XP magnetic beads (Beckman Coulter, #A63881), eluted in 12 µl of nuclease free water and quantify using the dsDNA QuantiFluor Dye System (Promega, #E2670).

The sequencing libraries were built by tagmentation using 50 ng of ds cDNA with Illumina Nextera XT Kit (Illumina, #FC-131-1024) following the manufacturer’s recommendations. The reaction was incubated 5 min at 55°C, immediately purified with DNA Clean & concentrator-5 Kit (Zymo research) and eluted with 21 µl of nuclease free water. Tagmented library was PCR amplified using 20 µl eluted cDNA, 2.5 µl of i7 Illumina Index, 2.5 µl of 5 µM P5-BRB primer (IDT) using the following program (72°C-3 min; 98°C-30 s; 13 cycles: 98°C-10 s, 63°C-30 s, 72°C-5 min). The fragments ranging 300-800 base pair (bp) were size-selected using SPRIselect (Bekman Coulter, #) (first round 0.65x beads, second 0.56x) with a final elution of 12 µl nuclease-free water. The resulting library was sequenced on Illumina Hiseq 4000 sequencer as Paired-End 100 base reads following Illumina’s instructions. Image analysis and base calling were performed using RTA 2.7.7 and bcl2fastq 2.17.1.14. Adapter dimer reads were removed using DimerRemover (https://sourceforge.net/projects/dimerremover/).

### BRB-seq raw data preprocessing

The first read contains 16 bases that must have a quality score higher than 10. The first 6 bp correspond to a unique sample-specific barcode and the following 10 bp to a unique molecular identifier (UMI). The second reads were aligned to the human reference transcriptome from the UCSC website (release hg38) using BWA version 0.7.4.4 with the non-default parameter “−l 24”. Reads mapping to several positions in the genome were filtered out from the analysis. The pipeline is described in Draskau (2021). After quality control and data preprocessing, a gene count matrix was generated by counting the number of unique UMIs associated with each gene (lines) for each sample (columns). The resulting UMI matrix was further normalized by using the rlog transformation implemented in the DeSeq2 package (Love et al., 2014).

### Bioinformatics analysis

Differential Gene Expression (DGE) analyses were performed using DeSeq2 R package (version 1.34.0, R 4.1) for each pairwise comparison of AB668, CX-4945 and DMSO conditions (Table S2). For these 3 DGE results, p-values were corrected by the Benjamini-Hochberg method. Only genes with corrected P-values < 0.05 and |Log2(Fold Change)|> 0.5 were considered as significantly differentially expressed. The first sample label in a comparison (e.g. AB668 for “AB668 vs DMSO”) means that it is the numerator in the calculation of the fold change (Fold Change = AB668/DMSO). The top 15 of significant genes based on FDR values and apoptotic and lipid & steroid metabolism related genes (if not overlapping with the top 15) are displayed in volcano plots. All DGE values for each selected genes in three comparisons can be retrieved in the supp-data (volcanoPlots3_AB668_CX_DMSO.ods). Gene Set Enrichment Analysis (GSEA) of Reactome pathway database (reactome.db version 1.82.0) was carried out using the Bioconductor/R package clusterProfiler (version 4.6.0). Pathways are considered significantly enriched whether related corrected P-value (Benjamini-Hochberg correction) is lower than 0.05.

## Acknowledgment

We thank Institut Convergence PLAsCAN (ANR-17-CONV-002), as well as ANR-21-CE18-0014-01 for project ANR-CK2COV, Ligue contre le Cancer (Comité de l’Isère), ITMO Cancer of Aviesan on funds administered by Inserm, Commissariat à l’Energie Atomique et aux Energies Alternatives (CEA), Université Grenoble-Alpes (UGA) and SATT PULSALYS for financial supports. We acknowledge both the European Union’s Horizon 2020 research and innovation program which funded P.B. and F.J. through the KATY project (grant agreement No 101017453) and the contribution of SFR Biosciences (UAR3444/CNRS, US8/Inserm, ENS de Lyon, UCBL): Protein Science Facilities. We are grateful to Dr David Knapp and Dr Andreas Krämer for the gift of the CK2α plasmid and their advice for CK2α crystallization. FJ and PB are supported by the KATY EU program Horizon 2020/H2020-SCI-FA-DTS-2020-1 (contract number 101017453).

We acknowledge the staff of the ID30B beamline at the ESRF. The CBS is a member of the French Infrastructure for Integrated Structural Biology (FRISBI), a national infrastructure supported by the French National Research Agency (ANR-10-INBS-05).

We thank Muriel Gelin for technical help for the treatment of the x-ray diffraction data.

