## Supplementary data for "AB668, a novel highly selective protein kinase CK2 inhibitor with a distinct anti-tumor mechanism as compared to CX-4945 and SGC-CK2-1"

**Figure S1. Binding sites and chemical structures of small-molecule CK2 inhibitors.** Compound 4 is a CK2 inhibitor that binds at the CK2α/CK2b interface; CX-4945 and SGC-CK2-1 are ATP-competitive inhibitors; CAM4066 and KN2 are bivalent inhibitors that bind at the ATP site and in the αD pocket of CK2. References and PDB codes are indicated for each molecule.

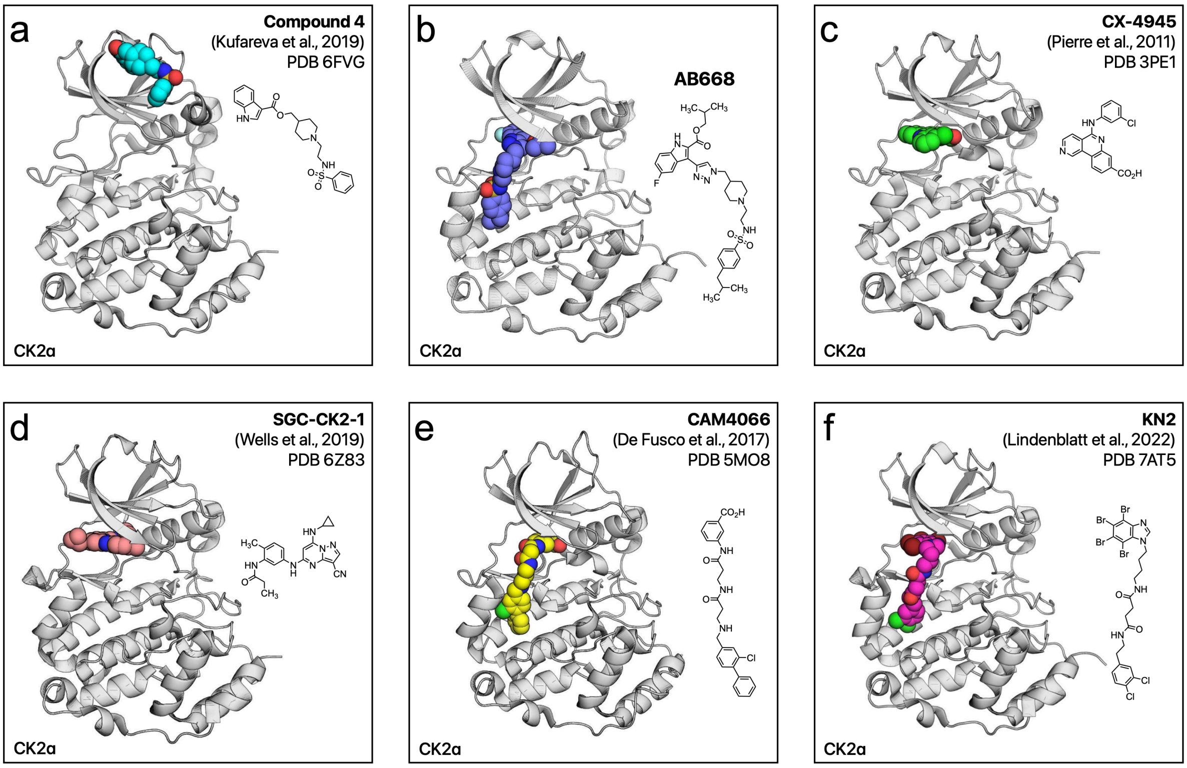

**Figure S2. OMIT electron-density-map for AB668 bound to CK2α.**
The purple mesh and surface represent the electron density map ($2F_O–F_C$ omit map) contoured at 1.0 σ of AB668 bound to CK2α.

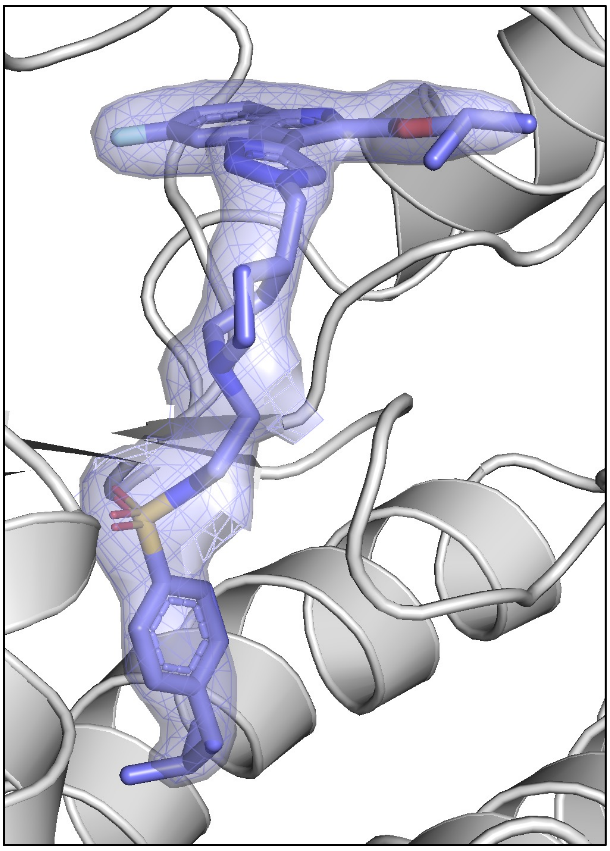

**Figure S3. Thermal Shift Assay experiments for the bivalent CK2 inhibitor AB668 and the ATP-competitive inhibitors CX-4945 and SGC-CK2-1 bound to CK2α.**
**a**) Thermal Shift Assay melting curves observed for CK2 alone (blue) and in the presence of the three inhibitors (25 mM). CK2α alone displayed a Tm of 43.6 ± 0.4°C. **b**) Variation of CK2α ΔTm upon inhibitor addition is reported as a function of the ligand concentration. At saturating inhibitor concentration, CX-4945 induced a ΔTm = 13.8 ± 0.4°C, SGC-CK2-1 a ΔTm = 11.0 ± 0.3°C and AB668 a ΔTm = 5.2 ± 0.4°C.

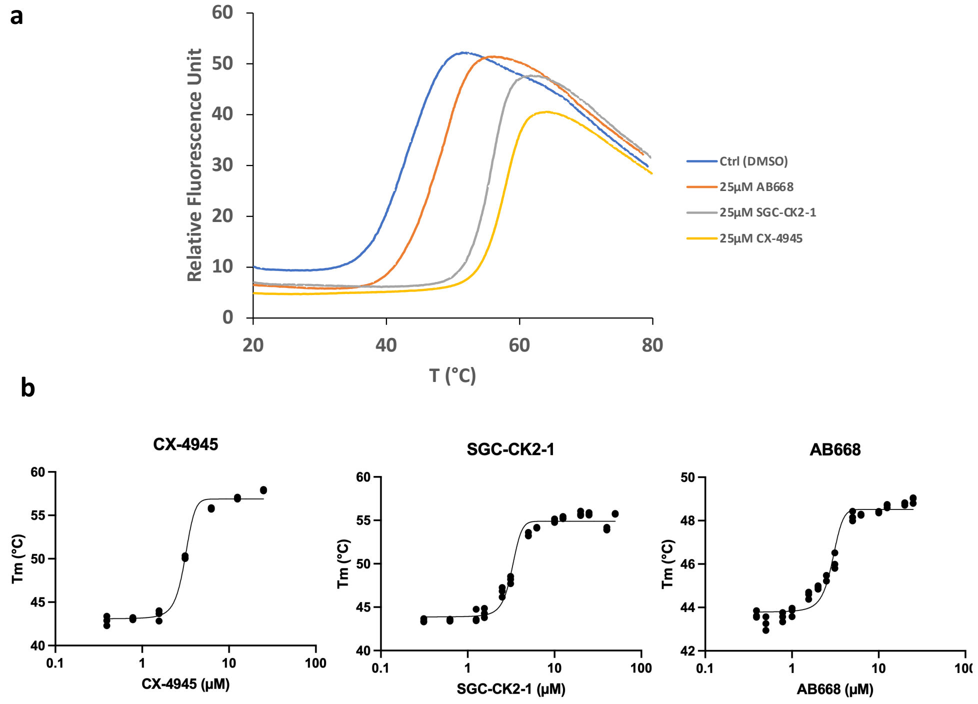

**Figure S4. Inhibition of CK2 by AB668.**

a) Inhibition constant for CK2 holoenzyme as determined using a canonical radiometric assay using a CK2β-dependent peptide substrate.

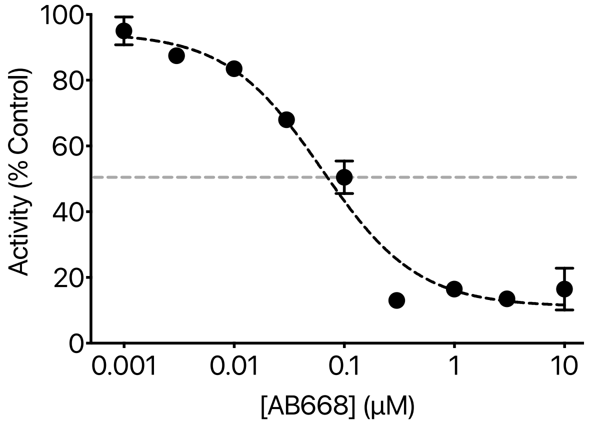

b) Inhibition of SIX1 by AB668. AB668 inhibits the phosphorylation of SIX1, a transcription factor that is phosphorylated by the holoenzyme. At 500 nM, AB668 inhibits about 90% of SIX1 phosphorylation

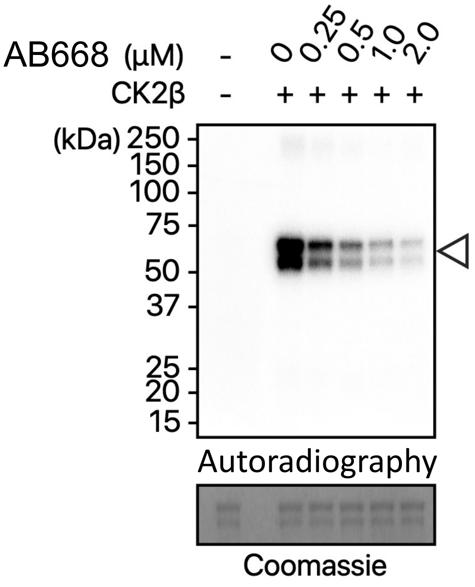

**Figure S5. Impact of AB668 in MDA-MB231 (Triple negative breast cancer) cells**
**a**) AB668 induces cell death in MDA-MB231 cells; **b**) normal MCF10A epithelial cells are insensitive to AB668; **c**) Western blot analysis in MDA-MB231 cells. p53 phosphorylation on Ser15 was only decreased at high AB668 concentrations, suggesting that p53 is not involved in MDA-MB231 cell death. p21 phosphorylation was also strongly downregulated by low concentrations of AB668.

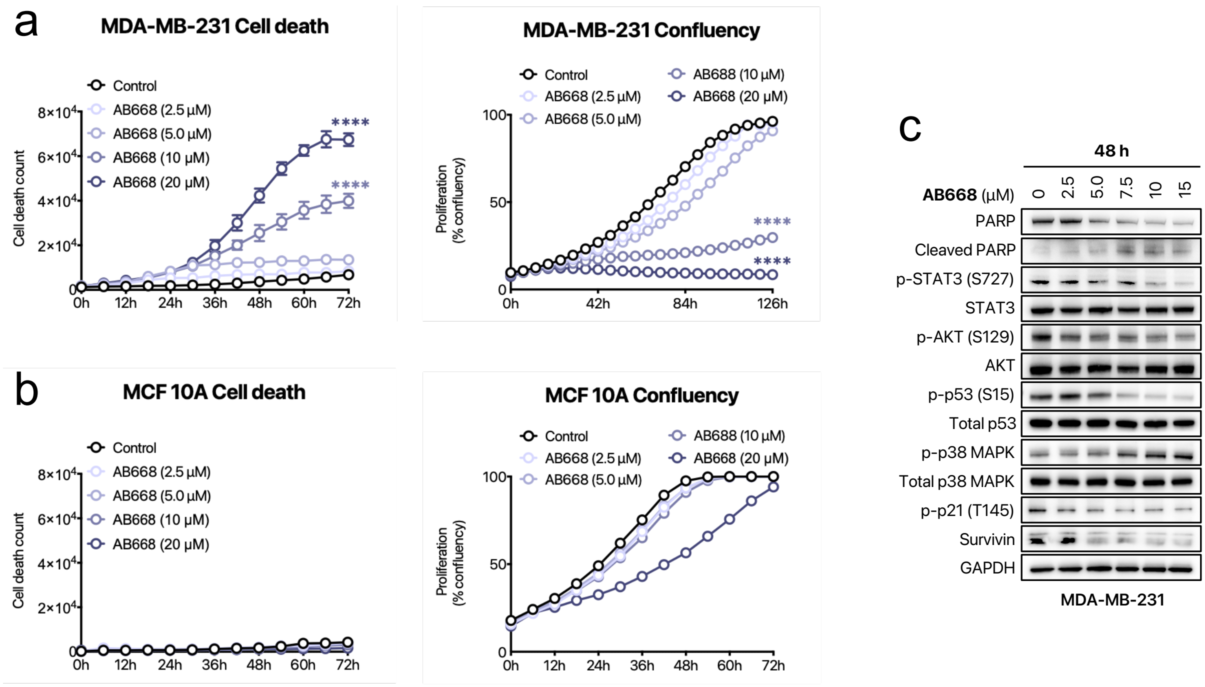

**Table S1. Data collection and refinement statistics (molecular replacement)**

| | AB668 |
|---|---|
| **Data collection** | |
| Wavelength (Å) | 0.97625 |
| Space group | $P43_{1}2$ |
| Cell dimensions | |
| $a, b, c$ (Å) | 126.785126.785 124.93 |
| $\alpha, \beta, \gamma$ (°) | 90.0, 90.0, 90.0 |
| Resolution (Å) | 51.64-2.50 (2.59-2.50) * |
| Total reflections | 197720 (19926) |
| Unique reflections | 35772 (3494) |
| Multiplicity | 5.5 (5.5) |
| Completeness (%) | 99.9 (100.0) |
| Mean $I / \sigma I$ | 8.9 (1.6) |
| Wilson B factor (Å$^2$) | 47.46 |
| $R_{merge}$ | 0.095 (0.768) |
| *Rmeas* | 0.105 (0.852) |
| *Rp.i.m* | 0.045 (0.359) |
| $CC_{1/2}$ | 0.984 (0.680) |
| Completeness (%) | 99.9 (100.0) |
| Redundancy | 5.5 (5.5) |
| **Refinement** | |
| $R_{work}$ / $R_{free}$ | 21.52/25.50 |
| No. atoms | |
| Protein | 5544 |
| Ligand | 179 |
| Water | 55 |
| $B$-factors | |
| Protein | 64.85 |
| Ligand | 72.40 |
| Water | 57.78 |
| R.m.s. deviations | |
| Bond lengths (Å) | 0.008 |
| Bond angles (°) | 0.972 |
| Ramachadran allowed | 5.79 |
| Ramachandran favoured | 93.45 |
| No. TLS group | 6 |

*Values in parentheses are for highest-resolution shell.

**Characterization of AB668**

AB888: isobutyl 5-fluoro-3-(1-((1-(2-((4-isobutylphenyl)sulfonamido)ethyl)piperidin-4-yl)methyl)-1*H*-1,2,3-triazol-4-yl)-1*H*-indole-2-carboxylate

**Mp** 161-163°C. **$^1$H NMR** (400 MHz, $d_6$-DMSO) δ = 12.01 (s, 1H), 8.55 (s, 1H), 8.05 (dd, *J* = 10.2, 2.5 Hz, 1H), 7.70 (d, *J* = 8.3 Hz, 2H), 7.55 (dd, *J* = 9.0, 4.7 Hz, 1H), 7.34 (t, *J* = 8.4 Hz, 3H), 7.22 (td, *J* = 9.1, 2.6 Hz, 1H), 4.33 (d, *J* = 7.0 Hz, 2H), 4.12 (d, *J* = 6.7 Hz, 2H), 2.86-2.78 (m, 2H), 2.65 (d, *J* = 11.2 Hz, 2H), 2.48 (s, 2H), 2.23 (t, *J* = 6.8 Hz, 2H), 2.03 (dq, *J* = 13.4, 6.7 Hz, 1H), 1.81 (tt, *J* = 16.5, 9.0 Hz, 4H), 1.44 (d, *J* = 11.3 Hz, 2H), 1.25-1.16 (m, 2H), 0.94 (d, *J* = 6.7 Hz, 6H), 0.81 (d, *J* = 6.6 Hz, 6H). **$^{13}$C NMR** (100 MHz, $d_6$-DMSO) δ = 161.0 ($C_{quat}$), 157.5 (d, *J* = 234.1 Hz, $C_{quat}$), 145.9 ($C_{quat}$), 140.2 ($C_{quat}$), 138.0 ($C_{quat}$), 133.1 ($C_{quat}$), 129.5 (CH), 126.4 (CH), 126.0 (d, *J* = 10.3 Hz, $C_{quat}$), 124.7 (CH), 123.7 ($C_{quat}$), 114.4 (d, *J* = 26.7 Hz, CH), 114.0 (d, *J* = 9.4 Hz, CH), 111.9 (d, *J* = 5.6 Hz, $C_{quat}$), 107.3 (d, *J* = 24.6 Hz, CH), 70.6 ($CH_2$), 56.8 ($CH_2$), 54.4 ($CH_2$), 52.5 ($CH_2$), 44.1 ($CH_2$), 40.2 ($CH_2$), 36.5 ($CH_2$), 29.5 (CH), 29.0 ($CH_2$), 27.4 (CH), 22.0 ($CH_3$), 18.9 ($CH_3$). **$^{19}$F NMR** (376 MHz, $d_6$-DMSO) δ = -122.4 (td, *J* = 9.7, 4.7 Hz). **HRMS (ESI$^+$) :** calcd. for $C_{33}H_{44}FN_6O_4S$, 639.3122. Found : $[M+H]^+$, 639.3123 (0.2 ppm error).

**$^{1}$H NMR** (400 MHz, $d_6$-DMSO) for compound AB668

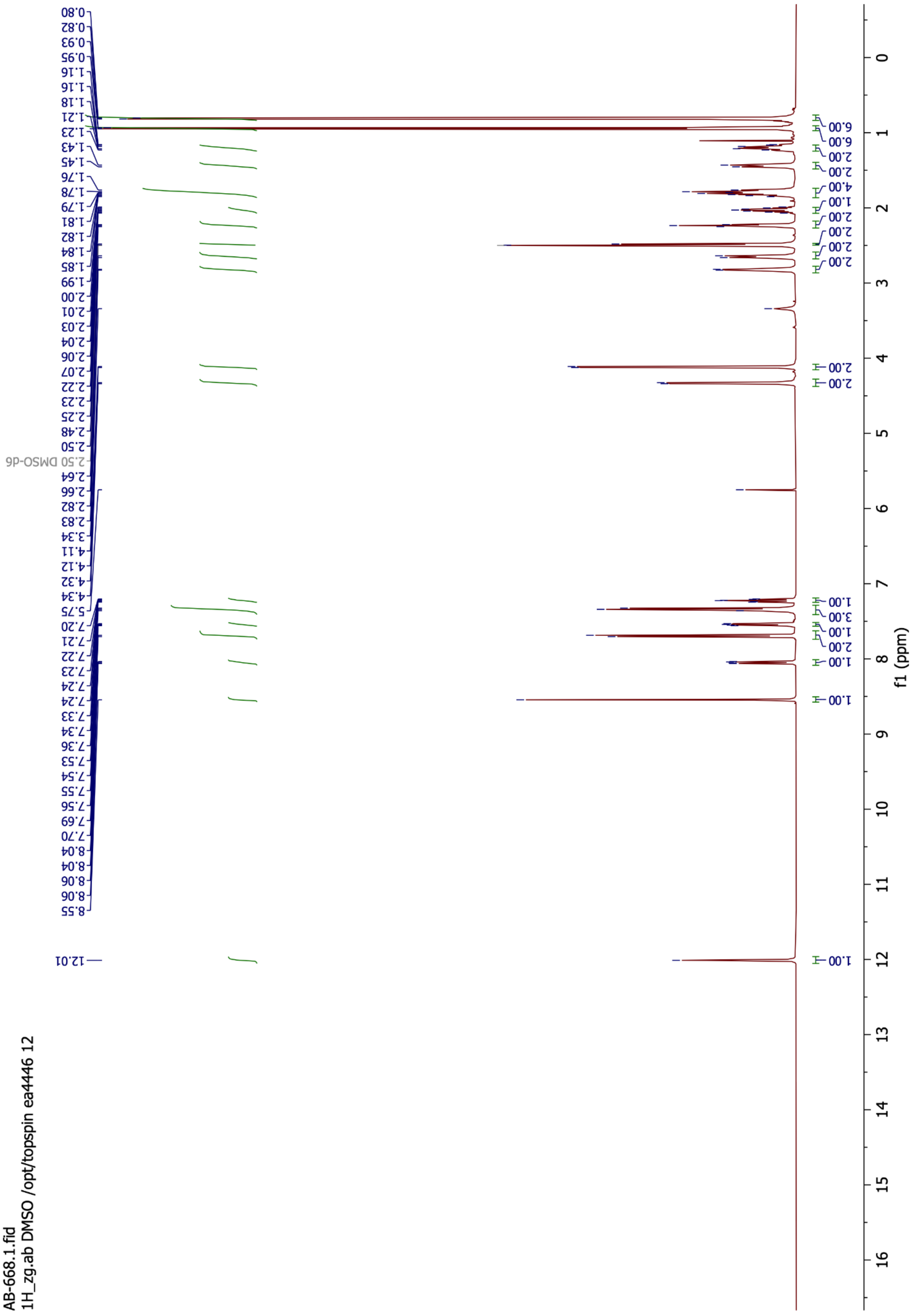

**$^{13}$C NMR** (100 MHz, d$_6$-DMSO) for compound AB668

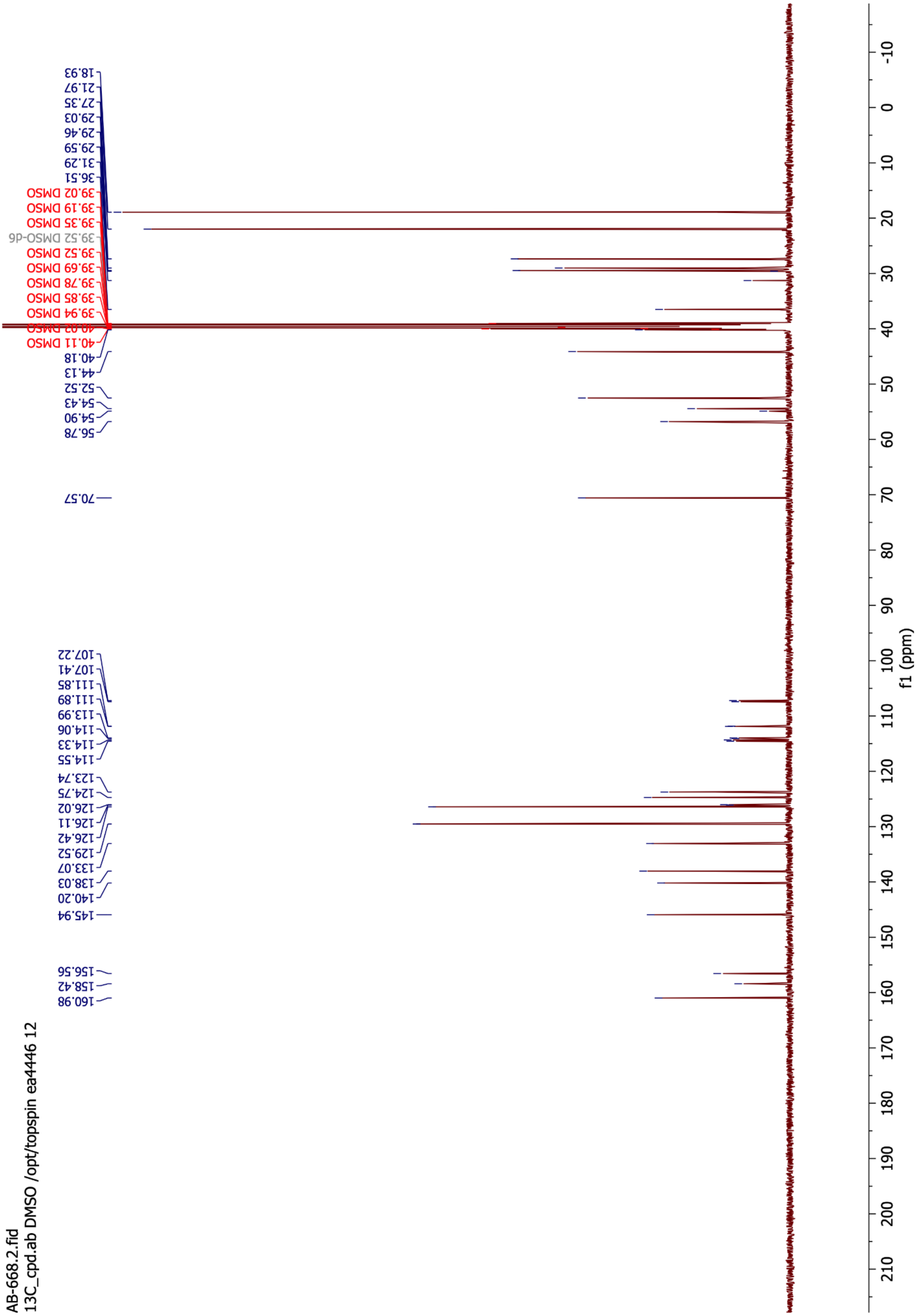

**$^{19}$F NMR** (376 MHz, $d_6$-DMSO) for compound AB668

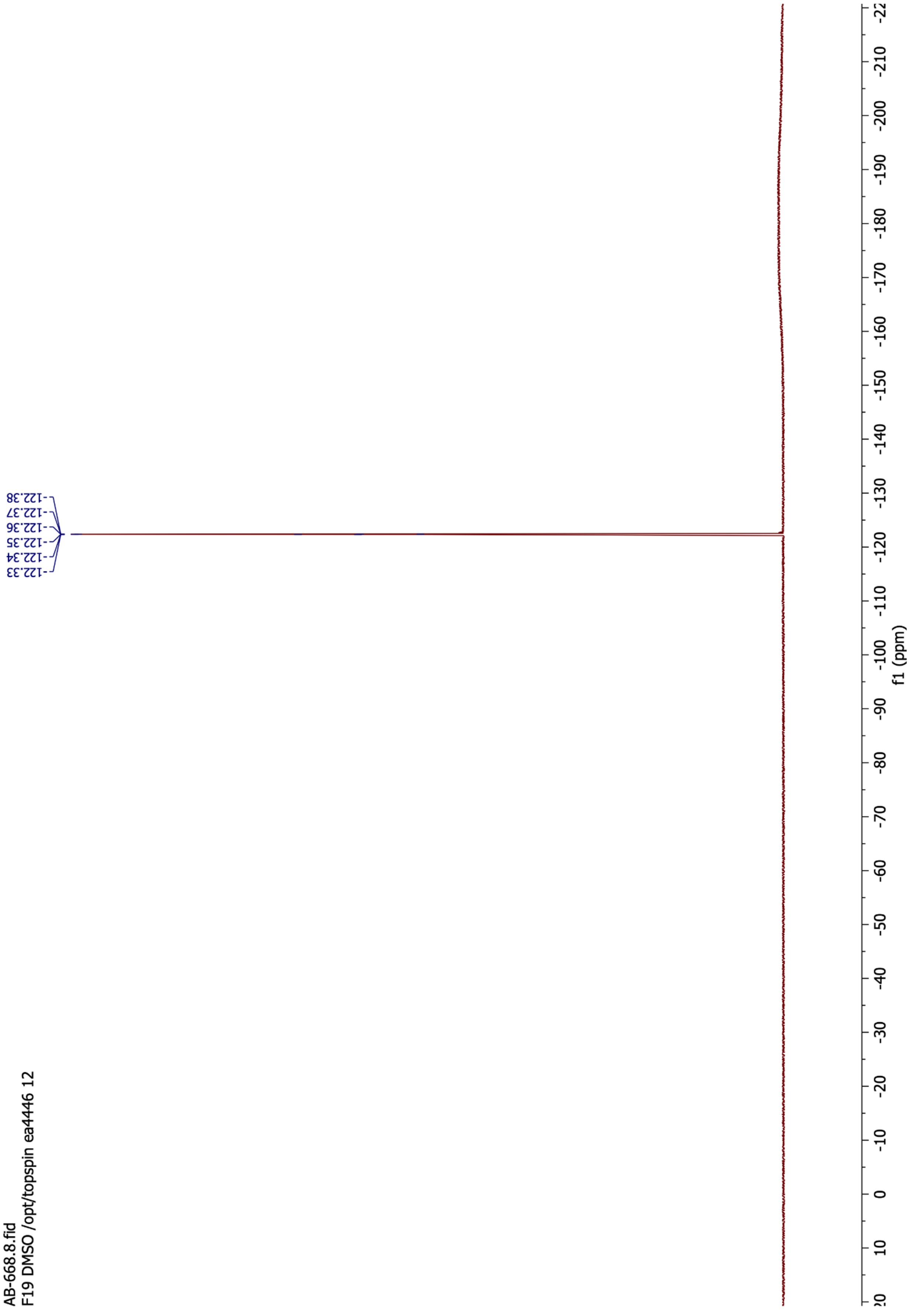
